# A novel network control model for identifying personalized driver genes in cancer

**DOI:** 10.1101/503565

**Authors:** Wei-Feng Guo, Shao-Wu Zhang, Tao Zeng, Yan Li, Jianxi Gao, Luonan Chen

**Affiliations:** Key Laboratory of Information Fusion Technology of Ministry of Education, School of Automation, Northwestern Polytechnical University, Xian, 710072, China; Key Laboratory of Systems Biology, Shanghai Institutes for Biological Science, Chinese Academy Science, Shanghai 200031, China; Department of Computer Science, Rensselaer Polytechnic Institute, Troy, NY, 12180, USA; Collaborative Research Center for Innovative Mathematical Modeling, Institute of Industrial Science, University of Tokyo, Tokyo 153-8505, Japan

**Keywords:** Driver genes, Structure network control theory, Tumor heterogeneity, Individual patients

## Abstract

Although existing computational models have identified many common driver genes, it remains challenging to identify the personalized driver genes by using samples of an individual patient. Recently, the methods of exploiting the structure-based control principles of complex networks provide new clues for identifying minimum number of driver nodes to drive the state transition of large-scale complex networks from an initial state to the desired state. However, the network control methods cannot be directly applied to identify the personalized driver genes due to the unknown network dynamics of the personalized system. Here we proposed the personalized network control model (PNC) to identify the personalized driver genes by employing the network control principle on genetic data of individual patients. In PNC model, we firstly presented a paired single sample network construction method to construct the personalized state transition network for capturing the phenotype transitions between healthy and disease state. Then, we designed a novel structure network control method from the Feedback Vertex Sets-based control perspective to identify the personalized driver genes. The experimental results on 13 kinds of cancer datasets from The Cancer Genome Atlas show that our PNC model outperforms other methods in terms of F-measures for identifying the personalized driver genes enriched in the gold-standard cancer driver gene lists. Thus PNC can provide novel insights for understanding tumor heterogeneity in individual patients.

**Author summary:** Notably there may be unique personalized driver genes for an individual patient in cancer. Identifying personalized driver genes that lead to cancer initiation and progression of individual patient is one of the biggest challenges in precision medicine. However, most methods for cancer-driver identification have focused mainly on the cohort information rather than on individual information and fail to identify personalized driver genes. We here proposed personalized network control model (PNC) to identify personalized driver genes by applying the network control principle on personalized data of individual patients. By considering the progression from the healthy state to the disease state as the network control problem, our PNC aims to detect a small number of personalized driver genes that are altered in response to input signals for triggering the state transition in individual patients. The impetus behind PNC contains two main respects. One is to design a paired single sample network construction method for constructing personalized state transition networks to capture the phenotypic transitions between normal and disease attractors. The other one is to develop a structure network control method on personalized state transition networks for identifying personalized driver genes which can drive individual patient system state from healthy state to disease state through oncogene activations. The experimental results on multiple cancer datasets highlight that PNC is effective for identifying personalized driver genes in cancer.

## Introduction

Cancer is a heterogeneous disease that is driven by oncogene activations such as genetic mutation, gene amplification, chromosomal rearrangement and transposable elements [1]. Researchers have recognized that personalized driver gene identification is required for discovering rare causal events in cancer and may provide vital information for selecting effective therapies in precision medicine[2, 3]. However, most computational models for cancer-driver identification look for genes that have more mutations than expected from the average background mutation rate. They mainly focus on cohort-level driver gene identification, and fail to identify the personalized driver genes. Therefore, new computational models and methods are urgently needed to identify personalized driver genes for understanding tumor heterogeneity in cancer. Recently, with the development of network science, structural-based network control approaches that aim to find a minimum set of driver nodes for steering the states of large-scale networks to the desired states can help the personalized driver genes identification in cancer propagation [2]. So far, the methods of exploiting the structure-based control principles of complex networks can be mainly divided into two categories according to the styles of the networks [4–7]. One focuses on the directed (or non-symmetrical) networks [4–6], and the other focuses on the undirected (or symmetrical) networks [7]. For directed networks, some linear or local nonlinear structural control tools based on Maximum Matching Sets (MMS) are developed to identify the minimum number of driver nodes that need to be controlled by external signals for the system to achieve the desired control objectives [4–6]. Although those MMS based control tools have many applications in modeling bio-molecular systems [2,9–11], those tools may only give an incomplete view of the network control properties for a system with nonlinear dynamics. Recently, the method of Feedback Vertex Set (FVS)-based control (FC) [12, 13] is proposed to control the large-scale networks in a reliable and nonlinear manner, where the network structure is prior known and the functional form of the governing equations is not specified, but it must satisfy some properties. With such a scheme, a Directed FVS-based control method (DFVS) is applied to control the directed networks by identifying the source nodes and FVS nodes as the driver nodes [8]. The above approaches only focus on the mainly linear or nonlinear dynamics of the directed networks. There are few approaches to investigate the structural controllability of the undirected networks. For example, the method of Minimum Dominating Sets (MDS) [7] investigates the structural controllability of the undirected networks. Since it works with the strong assumption that the controllers can control its outgoing links independently, MDS requires higher costs in many kinds of networks which may underestimate the structural control analysis of undirected networks. Therefore, efficient methods need to be developed for analyzing the structural controllability of the undirected networks.

Although these structure-based control methods of complex networks have offered powerful mathematical frameworks to understand diverse biological systems at a network level [2,9–11], they cannot be directly applied to the personalized driver genes identification of individual patients. This is primarily due to a gap between the structure network control methods and the applications in the personalized patient system. More efficient methods are needed to obtain personalized state transition networks that capture the phenotypic transitions between healthy and disease states, which is a rate-limiting step of structure network control methods. We addressed the limitation by introducing a novel personalized network control model (PNC), which sheds light on the cancer transition network between a disease state and a healthy state by employing the network control principle on the personalized genetic data of individual patients. PNC consists of the following two steps: (i) construct a personalized state transition network for characterizing the state transition of each individual during dynamical biological processes and (ii) design structure control method on the personalized state transition network. That is, we firstly developed a paired single sample network construction method (Paired-SSN) to construct the personalized state transition network for capturing the phenotype transitions between normal state and tumor state through the integration of the expression data and gene/protein interaction network. The personalized state transition network is a graph in which nodes represent genes, and edges denote the significant co-expression difference between normal state and tumor state in the gene interaction network. Then, on the personalized state transition networks, we designed a novel structure network control method (namely NCUA) according to the FC theory to drive a complex undirected network with nonlinear dynamics from initial attractor to desired attractor through perturbation to a feasible subset of driver nodes.The experimental results from 13 kinds of cancer datasets in The Cancer Genome Atlas (TCGA) demonstrate that:

i. Considering the Cancer Census Genes (CCG)[14] and Network of Cancer Genes (NCG) [15] as the gold-standard of cancer driver genes, PNC performs better in terms of the F-measures for identifying the personalized driver genes in contrast to traditional personalized driver genes identification methods (i.e., Differential expression genes identification methods and Hub genes selection method) and other personalized driver genes identification methods such as single sample controller strategy (SCS) [2]; furthermore, our PNC shows the enrichment significance in CCG and NCG on multiple cancer datasets;
ii. The F-measures of NCUA method for identifying the personalized driver genes enriched in CCG and NCG genes would be higher when Paired-SSN rather than LIONESS and SSN are used. When Paired-SSN are used for constructing the personalized state transition network, compared with other network control methods, NCUA can more effectively improve the performance in terms of the F-measures for identifying the personalized driver genes that are enriched in the CCG and NCG genes.
iii. For the cancer patient, the controllability and personalized driver genes evaluated by PNC vary in different cancer datasets; more heterogeneous the degree distribution of constructed personalized state transition network is, the easier it is to control the entire system. In conclusion, our PNC model provides a new powerful tool for identifying the personalized driver genes in individual patient system and gives novel insights for understanding tumor heterogeneity in cancer.

## Results

### An overview of PNC model

Cancer can be perceived as a dysfunction of molecular networks that regulate molecular communications and cellular processes [2–3]. To fully understand cancer progression, we need to understand the dynamics of these networks with respect to control theory [16]. In particular, we consider the progression from a healthy state to a disease state as a network control problem. By considering the gene expression profiles in normal and tumor samples as the respective state for a given patient and applying adequate knowledge of the network structure, structure network control tools aim to detect a small number of genes that are altered in response to input signals and trigger a state transition in individual patients. The personalized driver genes are the altered nodes/genes in response to input signals which can trigger the state transition of the whole biological network. The input signals may be oncogene activation signals such as a genetic mutation, gene amplification, a chromosomal rearrangement, or transposable elements. The “controllers” in PNC model for identifying personalized driver genes are the genetic or environment factors which produce the oncogene activation signals.

Our PNC consists of two main parts. One is a paired single sample network construction method (Paired-SSN) rather than an original SSN [17], which we used to construct personalized state transition networks. In these networks, edges exist in both the gene interaction network and the differential co-expression network for each patient based on integration of the matched genetic information of normal and disease samples from the same patient (Figure 1). In this work, we used a gene interaction network from literature [18] by integrating multiple types of datasets, including Mutual Exclusivity Modules in cancer [19, 20], Reactome [21], NCI-Nature Curated PID [22] and Kyoto Encyclopedia of Genes and Genomes[23]. The gene interaction network consists of 11,648 genes and 211,794 interactions. The nodes in the personalized state transition networks represent the genes, and a pair of nodes denotes the significant co-expression difference of gene intaractions between normal sample and tumor sample. For the second part of PNC, we designed a novel nonlinear structural control method (namely, NCUA) [12, 13] for identifying personalized driver genes. This was done to ensure that the system state would asymptotically be changed in the personalized state transition network. These two primary methodologies are described in more detail in the section of Methods.

**Figure 1.**
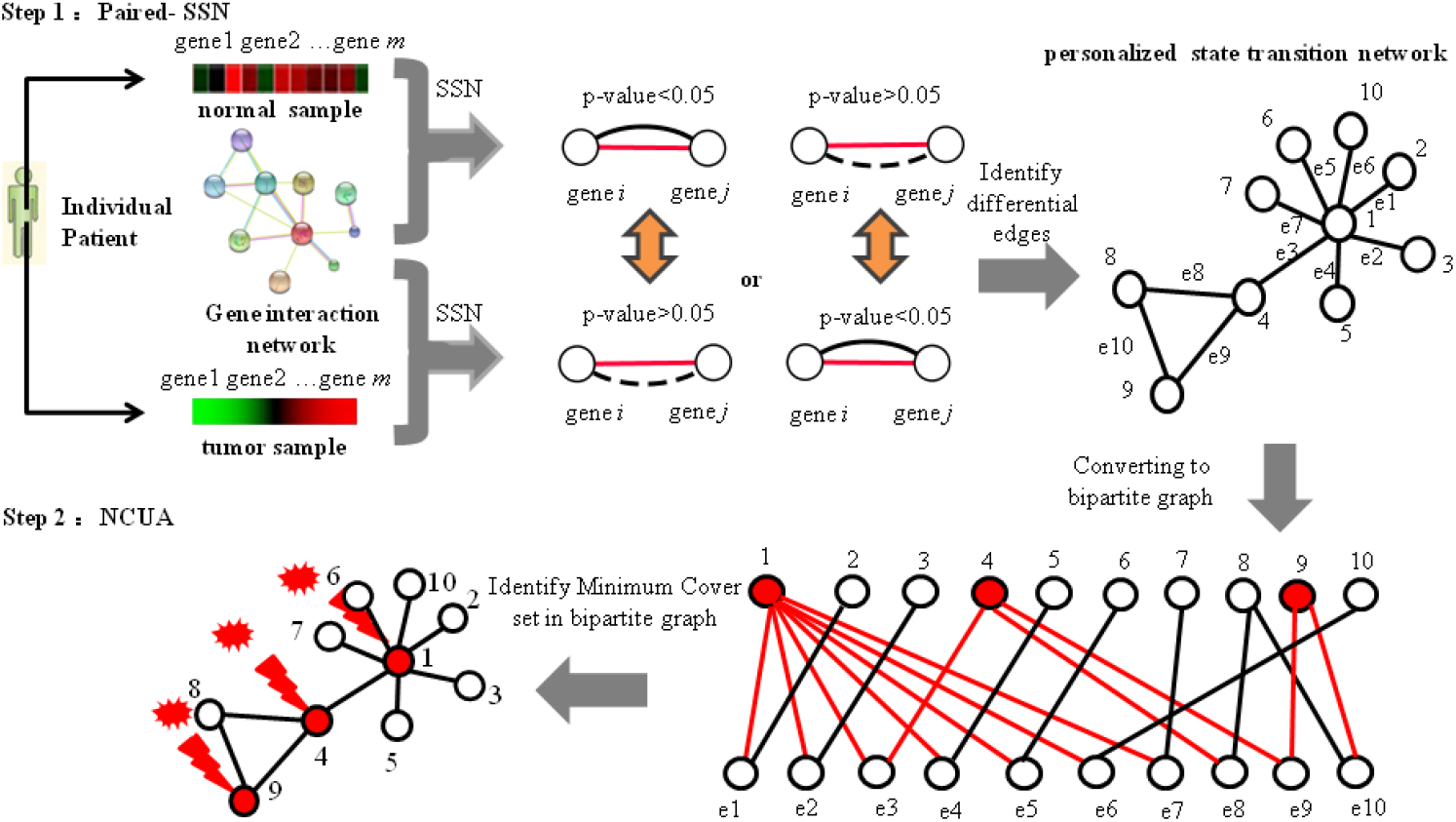
Overview of personalized network control model (PNC) for identifying the personalized driver genes. PNC model consists of two main parts. One is that a Paired-SSN method was developed to construct the personalized state transition networks by using the matched expression information of normal and tumor samples of an individual patient and gene interaction network. For an individual patient, we firstly respectively obtained co-expression p-value of the interaction edges for normal sample and tumor sample by using the SSN. Then the personalized state transition network was constructed. In the personalized state transition network, the edge between gene *i* and gene *j* will exist if co-expression p-value of the interaction edge is less than (greater than) 0.05 for the tumor sample but greater than (less than) 0.05 for the normal sample. The solid line denotes that co-expression p-value is less than 0.05 while the dotted line denotes that co-expression p-value is larger than 0.05. The red solid line denotes the interaction edge in gene interaction network. Therefore edges in the personalized state transition networks denote the significant co-expression difference of gene intaractions between normal sample and tumor sample. Another part is to design a novel nonlinear structural control method (NCUA) for identifying the personalized driver genes by considering personalized state transition network as an undirected network. For our NCUA method, we assumed that each edge in an undirected network is bi-directional (feedback loop) and constructed a bipartite graph from the undirected network. In bipartite graph, the nodes of top side are the nodes of original graph and the nodes of the bottom side are the edges of the original graph. In the end, we adopted an equivalent optimization procedure for obtaining the minimum dominating nodes in the up side to cover the bottom side nodes in the bipartite graph, which are considered as the minimum driver nodes of NCUA.

### Performance comparisons of PNC with other personalized driver genes identification methods

To identify personalized driver genes by using PNC, we collected paired samples (a control/normal sample and a tumor sample) from each individual. Here we used cancer datasets which contained enough normal-disease paired samples (>20 paired samples) in TCGA for the case study. By searching TCGA, 13 cancer datasets met the requirements, i.e., the datasets for breast invasive carcinoma (BRCA), colon adenocarcinoma (COAD), kidney chromophobe (KICH) and kidney renal clear cell carcinoma (KIRC), kidney renal papillary cell carcinoma (KIRP), liver hepatocellular carcinoma (LIHC), lung adenocarcinoma (LUAD), lung squamous cell carcinoma (LUSC), stomach adenocarcinoma (STAD), uterine corpus endometrial carcinoma(UCEC), Head and Neck Squamous Cell Carcinoma (HNSC),Prostate Adenocarcinoma (PRAD) and Thyroid Papillary Carcinoma (THCA). More information is given in Table S1 of Additional File 1.

On the 13 cancer datasets, the identified personalized driver genes annotated in the Cancer Census Genes (CCG)[14] and Network of Cancer Genes (NCG) [15] (Additional File 2) were adopted to compute the F-measure scores (Methods) for assessing the performance of PNC. Figure 2(a) is the results of PNC and other traditional methods such as DEG-FoldChange, DEG-p-value, DEG-FDR and Hub genes selection methods (Methods). DEG-FoldChange selects the personalized driver genes by calculating the fold-change between normal sample and tumor sample (|log2(fold-change)|>1). DEG-p-value and DEG-FDR selects the personalized driver genes by respectively calculating the p-value and FDR (<0.05) between a cancer tumor sample and a group of control samples. Hub genes selection method regards the hub genes in the constructed network as cancer driver genes. From Figure 2(a), we can find that F-measures of PNC are higher than that of DEG methods (i.e., DEG-FoldChange, DEG-p-value and DEG-FDR) and Hub genes selection method. These results show that our network control method is more effective for discovering personalized driver genes than the DEG methods and the Hub genes selection method.

**Figure 2.**
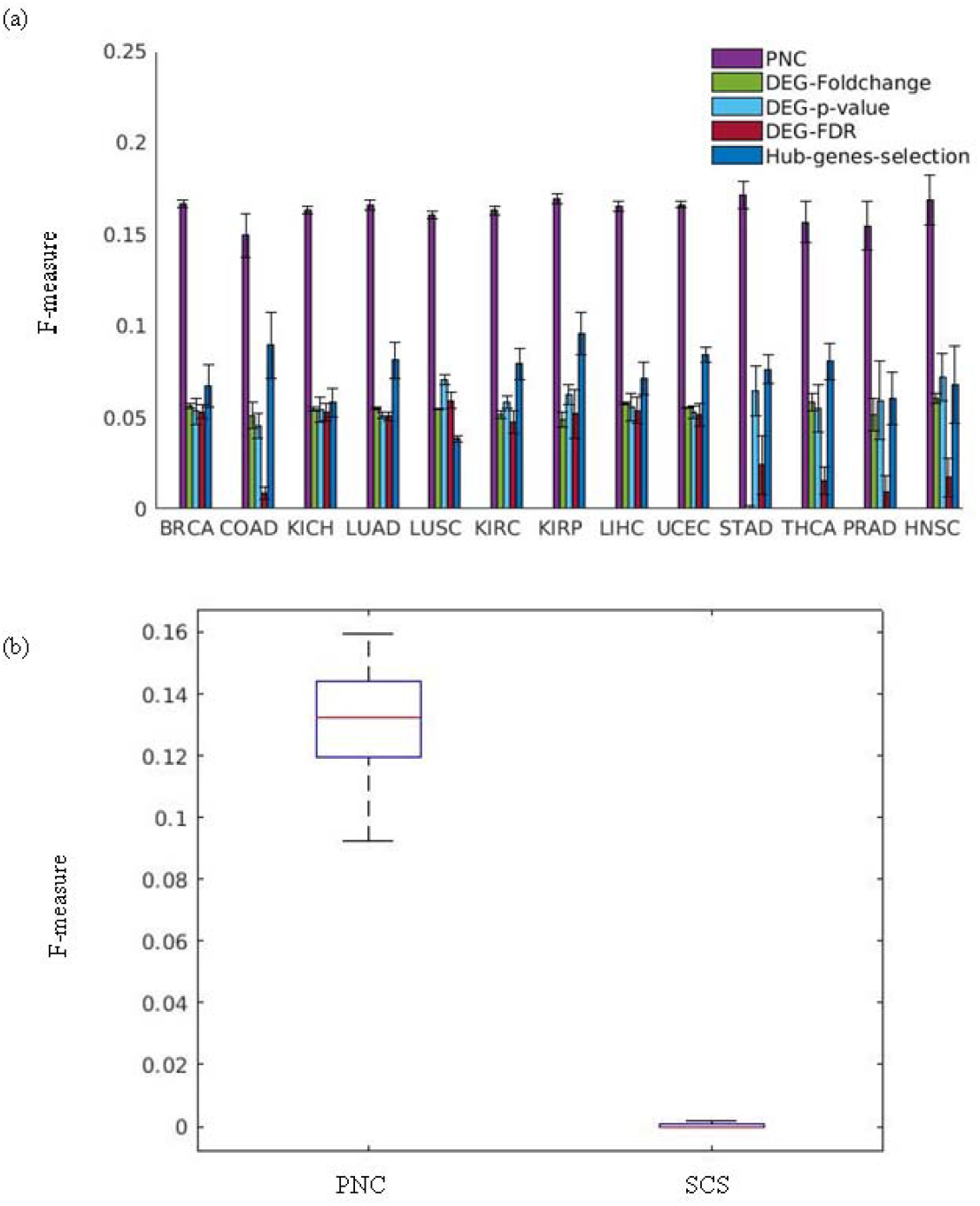
Results of PNC and other methods. (a) F-scores of PNC and other four traditional methods (i.e., DEG-FoldChange, DEG-p-value, DEG-FDR and Hub genes selection method) on 13 cancer datasets. (b) F-scores of PNC and single sample controller strategy (SCS) [2] on the PRAD cancer dataset.

Recently a single-sample controller strategy (SCS) based on the MMS structural control theory [2] was proposed to identify personalized driver genes. SCS integrates mutation data (e.g., single nucleotide variations, SNV; copy number variations, CNV) and expression data (i.e., paired samples of each patient) into a reference molecular network for each patient to obtain the driver genes in a personalized-sample manner. To further evaluate the advantage of PNC, we obtained the single nucleotide variations mutation data, the copy number variations mutation data and the gene expression data of the PRAD cancer dataset and compared the performance of PNC with that of SCS in terms of F-measures for identifying personalized driver genes enriched in CCG and NCG genes. Figure 2(b) shows the comparison results of PNC and SCS. We can see that PNC has higher F-measure than SCS, indicating that PNC can more effectively identify personalized driver genes which are enriched in the CCG and NCG genes.

### The effect of the different single sample network construction methods

In addition, to evaluate the influence of potential different personalized state transition network construction methods, we used LIONESS and SSN for constructing personalized state transition networks on the collected 13 cancer datasets. Different from Paired-SSN, SSN considers the tumor sample network as the personalized state transition network. The tumor sample network is constructed by quantifying the differential network between the tumor sample and a group of normal samples. LIONESS[24] constructs the personalized state transition networks by calculating the edge statistical significance between all tumor samples and the tumor samples without a given single sample (Methods). As LIONESS can be applied to multiple aggregate network reconstruction approaches, we used the Pearson Correlation Coefficient (PCC) in LIONESS to guarantee a fair comparison with the SSN and Paired-SSN methods. To keep the edge direction and filter the noise of PCC correlation in the personalized state transition networks, the previously mentioned gene interaction network in Paired-SSN was used on the LIONESS and SSN.

Based on the personalized state transition networks constructed by LIONESS, SSN and Paired-SSN respectively, we applied the NCUA method for identifying the personalized driver genes (Figure 3 (a)). From Figure 3 (a), we found that considering the CCG and NCG genes as the gold-standard of cancer driver genes, the F-measures of network control methods would be higher when SSN and Paired-SSN networks rather than LIONESS were used. Furthermore, F-measures of NCUA on Paired-SSN are more robust than SSN method. For example, in the KIRP cancer dataset, NCUA on Paired-SSN had higher F-measure than SSN method [17] although NCUA on Paired-SSN and SSN exhibited a similar performance in other cancer datasets. Therefore compared with other methods, Paired-SSN can include more information of individual patients and be suitable for identifying the personalized driver genes of individual cancer patients in this work.

**Figure 3.**
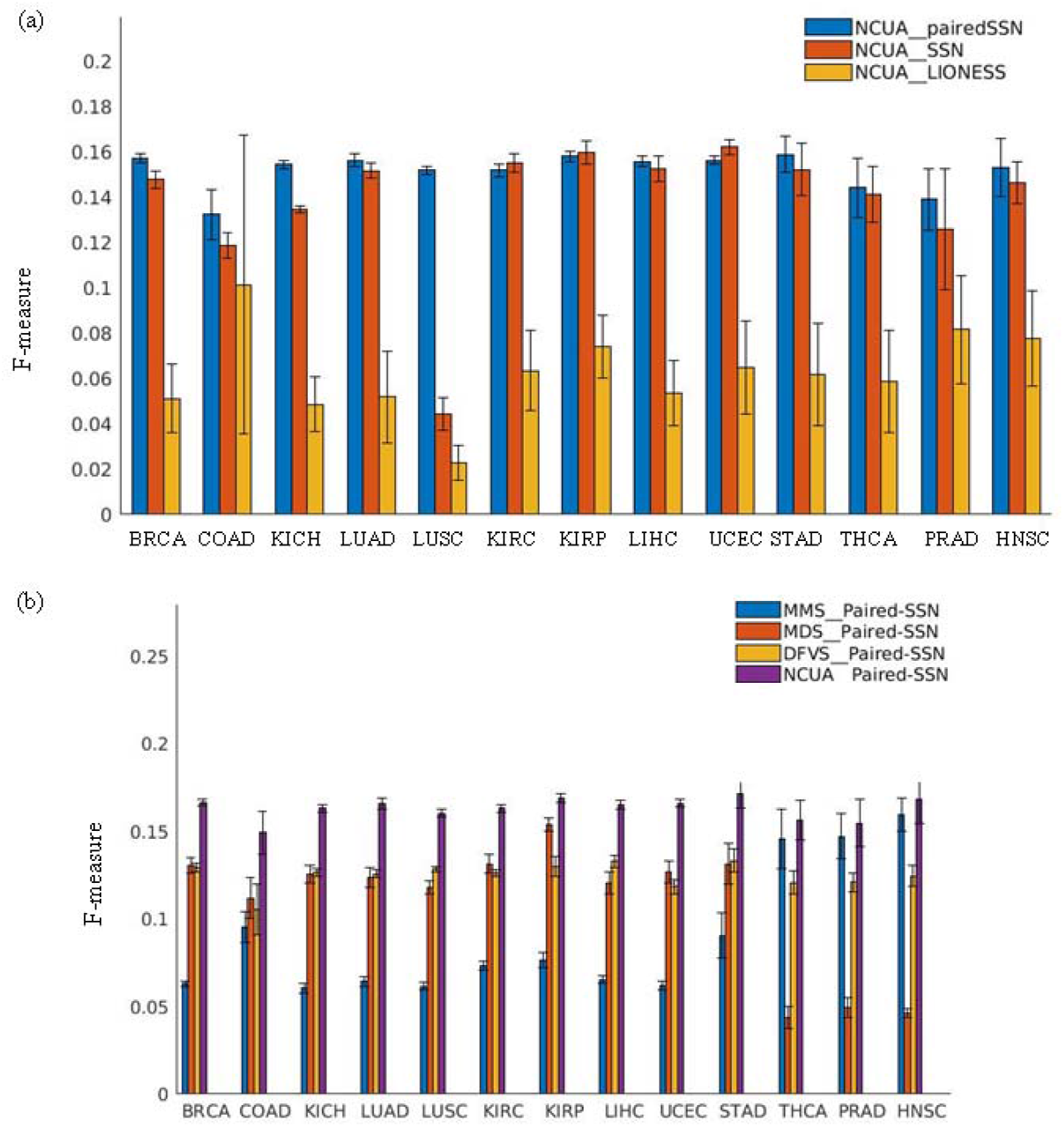
The effect of different single sample network control methods and network control methods on PNC. (a) The significant enrichment F-scores of NCUA on personalized state transition networks constructed by Paired-SSN (the first step of our PNC), SSN and LIONESS. (b)The enrichment F-scores of NCUA (the second step of our PNC) and other structure network control methods (MMS and MDS and DFVS) on the paired SSN networks (the first step of our PNC) in the list of CCG and NCG genes.

### The effect of the different network control methods

Furthermore, to evaluate the efficiency of NCUA as the second step of PNC, we gave computational comparisons in terms of F-measures for identifying personalized driver genes enriched in CCG and NCG genes between our NCUA and other methods including MMS (full control) [4], MDS[7] and DFVS[8] (Figure 3 (b)). We should note that MMS and DFVS are directed-network-based control methods while MDS and NCUA are undirected-network-based control methods. Thus, for MMS and DFVS, the personalized state transition network is a directed graph in which the directed information of reference gene interaction is used. Meanwhile, for MDS and NCUA, the personalized state transition network is an undirected graph in which the directed information of reference gene interaction is not used. In Table S2 of Additional file 1, we gave the concept comparisons including the network types and targeted states and time complexity and network dynamics between our NCUA and other network control method including MMS, MDS and DFVS. From Figure 3 (b), we can find that F-measures of NCUA in 13 cancer datasets are higher than those of MMS, MDS and DFVS methods, so our NCUA could overall identify personalized driver genes more efficiently than other network control methods.

### Statistic analysis of the personalized driver genes identified by PNC

In fact, PNC provides a set of personalized driver genes for each patient. On each cancer dataset, by calculating the frequency of genes that appear as the personalized driver genes among a group of patients, we defined high-frequency personalized driver genes (*fh*>0.6), medium-frequency personalized driver genes (0.3<*fm*≤ 0.6), and low-frequency personalized driver genes (*fl*≤ 0.3). In Figure 4, we gave the fractions of high-frequency personalized driver genes (yellow), medium-frequency personalized driver genes (red), and low-frequency personalized driver genes (blue) on 13 cancer datasets. Figure 4 revealed that different cancer datasets had different distributions for personalized driver genes among groups of patients. For example, in the PRAD cancer dataset, the high-frequency personalized driver genes accounted for the majority (>60%); however in the UCEC cancer dataset, the high-frequency personalized driver genes accounted for a minority (<20%). These results demonstrated that the heterogeneity of personalized driver genes varied in different cancer datasets.

**Figure 4.**
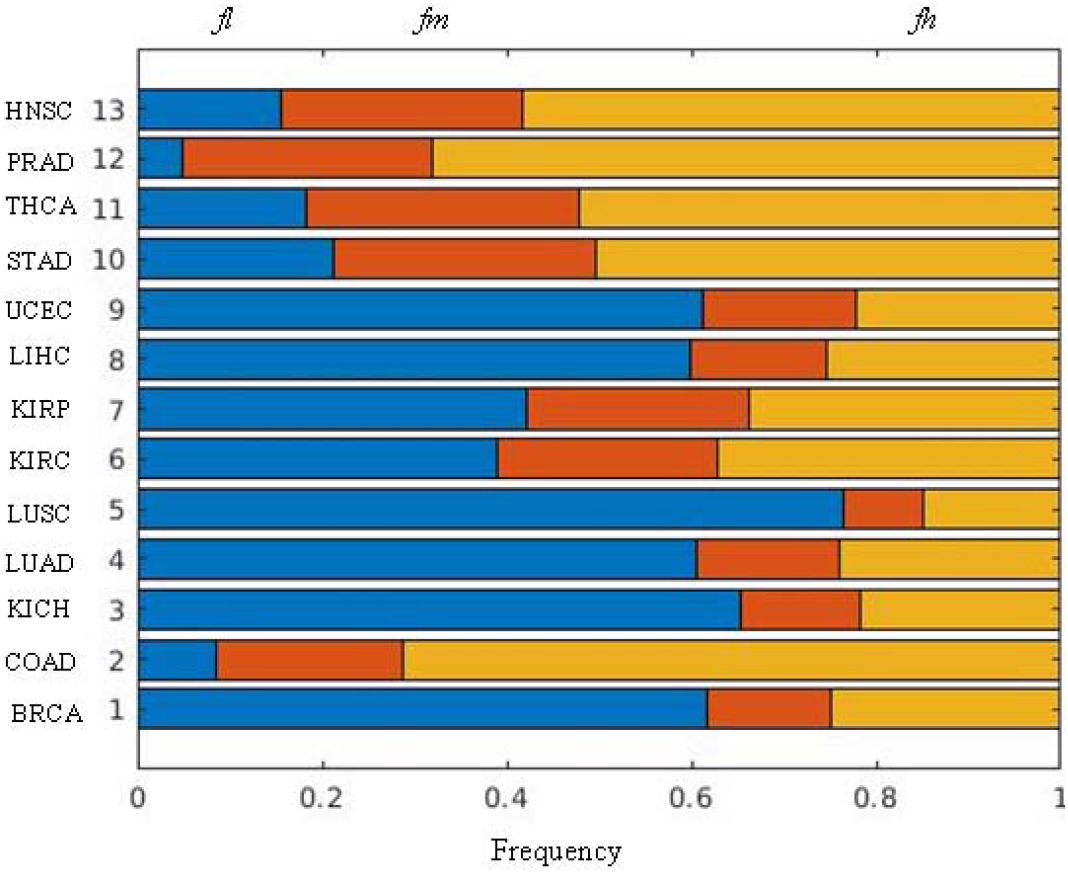
The fractions of high-frequency personalized driver genes (yellow, *fh*), medium-frequency personalized driver genes (red, *fm*), and low-frequency personalized driver genes (blue, *fl*) on 13 cancer datasets.

It is widely accepted that driver genes have low mutation frequency, which is called “long tail of rarely mutated genes” [25]. To demonstrate the ability of discovering the driver genes with rare mutation frequency, we first obtained the single nucleotide variations (SNVs) data of 13 kinds of cancer datasets from TCGA [26]. We then divided the personalized driver genes within CCG and NCG genes into driver genes with rare mutation frequency (0<frequency<0.05) and high mutation frequency (frequency>0.05) respectively. Finally we calculated the fraction of personalized driver genes with rare mutation frequency and high mutation frequency respectively on 13 kinds of cancer data as shown in Figure 5. Results in Figure 5 show that our PNC tends to discover personalized driver genes with rare mutation frequency on most of cancer datasets. However the fraction of the driver genes with rare mutation frequency and high mutation frequency are similar on UCEC and STAD cancer datasets.

**Figure 5.**
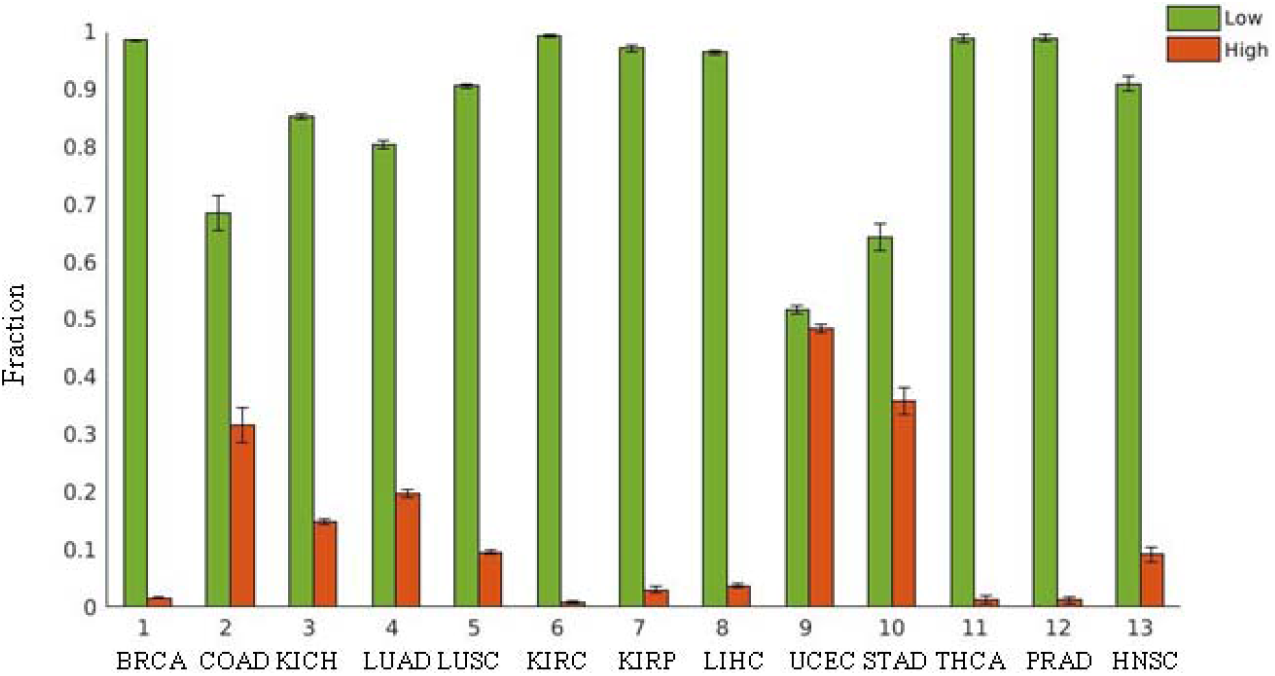
The fraction of personalized driver genes in CCG and NCG with low mutation frequency and high mutation frequency respectively on 13 kinds of cancer data.

### The enrichment analysis of the personalized driver genes identified by PNC

To further support the efficiency of PNC by statistic significance, the enrichment p-values of the predicted driver genes in CCG and NCG lists were evaluated for PNC as shown in Figure 6. The enrichment p-values were calculated by using the hyper geometric test [27]. The computational details were shown in section of Methods. From the results of Figure 6, we can see that PNC is actually significant for predicting the personalized driver genes enriched in the CCG and NCG lists. Furthermore, from Figure 6, we can see the following facts, for the CCG and NCG lists, the enrichment p-values of PNC vary in the different cancer datasets. For example, for COAD, PRAD, THCA and HNSC cancer dataset, not all patients have significant enrichment results (Figure 6). But for other cancer datasets such as BRCA, KICH, LUAD and UCEC cancer datasets, the personalized driver genes are significant enriched in CCG and NCG genes (p-value<0.05) for all patients by using the PNC method (Figure 6).

**Figure 6.**
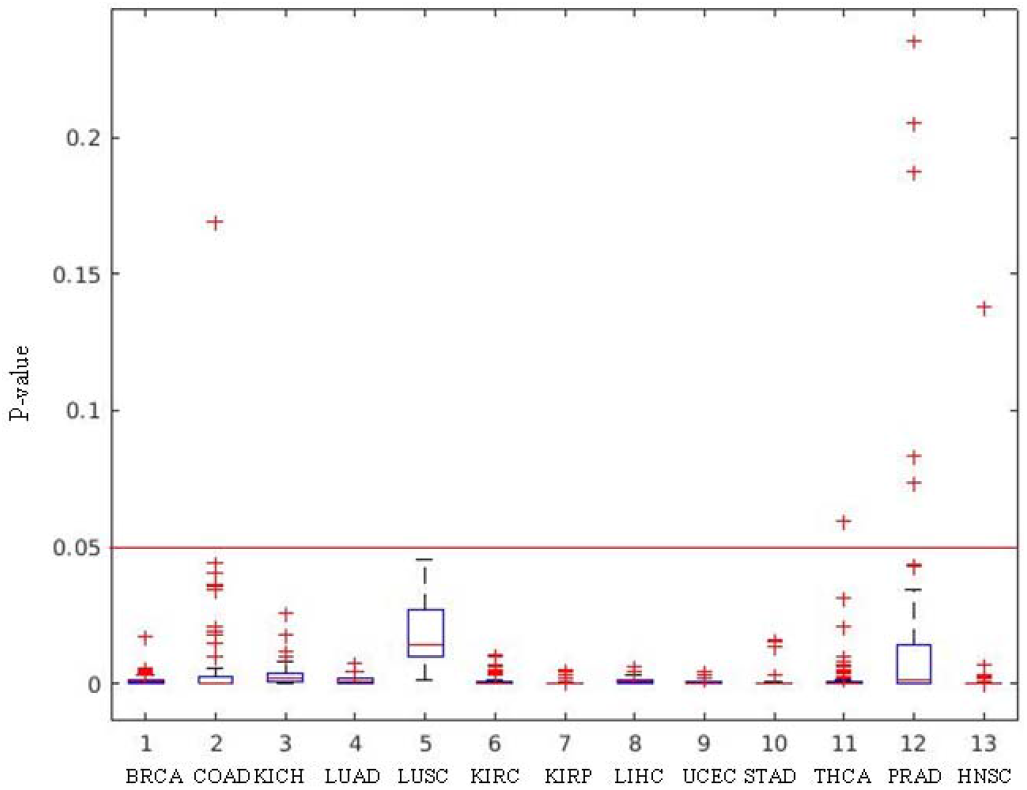
The enrichment p-value of personalized driver genes on 13 kinds of cancer data. The red line denotes the significant threshold value 0.05.

### Controllability evaluation of individual patient system by using PNC

To discover the personalized controllability of individual patient system, we employed the PNC on 13 cancer datasets in TCGA. We defined controllability as follows,

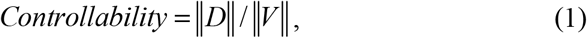

where ∥*D*∥ denotes the set of driver nodes to control the whole network and ∥*V*∥ denotes the number of connected nodes in the network. In Eq. (1), the smaller the value of *controllability* is, the more easily the personalized state transition network will be controlled. The personalized controllability of PNC on 13 cancer datasets from TCGA are shown in Figure 7 (a). From Figure 7 (a), we can find that the personalized controllability of individual patients varies in different cancer datasets. For example, the personalized controllability in COAD cancer data is less than 0.4, while the personalized controllability in LUSC is higher than 0.5.

**Figure 7.**
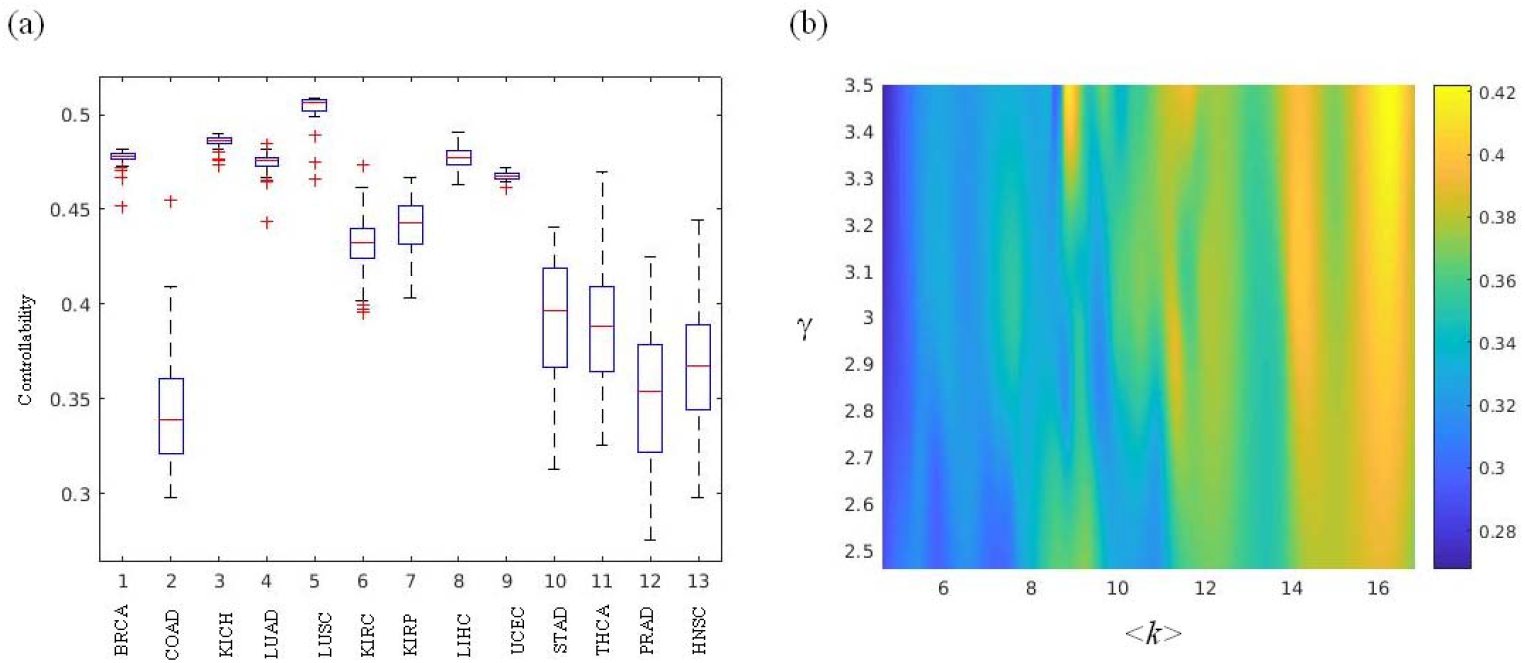
Controllability evaluation of individual patient system by using PNC. **(a)** Box plot with the distribution of the personalized controllability in 13 cancer datasets. **(b)** Heat map of the personalized controllability (different colors) which is as a function of the average degree and power law degree exponent.

To demonstrate the relationship between personalized controllability and network structure, we firstly obtained the connected component (CC) of each personalized state transition network and then used the CC network to determine the degree exponent γ by using the Kolmogorov-Smirnov goodness-of-fit test [28]. More computational details are shown in Supplementary Note 4 of Additional file 1. The network parameters in each CC on different cancer datasets including number of nodes, average degree and degree exponent are presented in Additional file 3. Finally in Figure 7 (b) we gave the controllability (defined in Eq. (1)) of personalized state transition networks which are significantly subject to the power-law distribution (p-value>0.05) as a function of the average degree and power law degree exponent. From Figure 7 (b), we found that the personalized state transition networks with smaller values of the average degree and power law degree exponent would be easier to control. That is, the more heterogeneous the degree distribution of a personalized state transition network is, the easier it is to control the entire system.

To more clearly explain how network parameters affect the controllability, we applied NCUA on multiple synthetic scale-free (SF) networks [29], as shown in Figure S1 of Additional file 1. Figure S1 show that the more heterogeneous an SF network degree distribution is, the easier it is to control the entire system. Those simulation results match well with the results of experimental results in cancer. Furthermore when the degree exponent γ is above 3, the fraction of minimum driver nodes is larger than that of the Erdös-Rényi random networks (Figure S1). These results gave insight into heterogeneous networks will be easier to control with the minimum number of driver nodes than the homogeneous networks. More details are shown in supplementary note 3 and Supplementary Note 5 of Additional file 1.

## Discussion

Cancer is known as a disease mainly caused by gene alterations [30]. Identification of driver genes is a key point of focus in cancer genomics. Computational models and methods are required to prioritize biologically efficient driver genes dependent on cancer high-throughput sequencing data. However, few methods can efficiently distinguish the personalized driver genes that change the state of genes in each patient. Recently, structure-based network control approaches have enabled us to investigate how to control complex networks by using a minimum set of driver nodes which could enable us to understand mechanisms of cancer progression. For structure-based network control methods, a minimum number of driver nodes need to be identified to drive the network state from the initial state to the desired state based on adequate knowledge of the network structure. The application of network control methods for personalized driver genes identification requires two key steps, i.e., (i) to construct personalized state transition networks which are involved in the state transition during disease development for each patient and (ii) to design the optimal network control methods based on the structure of the personalized state transition network. Here, we propose PNC based structure network control theory to personalized driver genes identification. PNC considers how to construct the personalized state transition network capturing the phenotype transitions between healthy and disease state through the utilize of Paired-SSN method and find the personalized driver genes on phenotype transitions by applying NCUA, which can control the individual from the normal attractor to the disease attractor.

To further demonstrate the advantage of our PNC, it has been employed to investigate personalized driver genes of cancer samples from TCGA by screening known driver genes in the CCG and NCG genes as controls of their phenotype transitions. We find that in contrast to other models on the personalized driver genes identification such as Differential expression genes identification methods, Hub genes selection method and single sample controller strategy (SCS), our PNC shows a higher performance in identifying the personalized driver genes in the CCG and NCG genes. Particularly we have validated that Paired-SSN has better performance for constructing personalized state transition networks compared with other methods, such as SSN and LIONESS. Furthermore NCUA is a better choice for the identification of personalized driver genes on the constructed personalzied state transition networks compared with other network control methods. Although NCUA already performs better than other network control methods, efforts to achieve promising improvements for the utilization of the NCUA approach are still ongoing. For example, NCUA does not consider multiple driver-node sets to control the network. Therefore in the future it is worth studying how to find multiple driver nodes configurations in large-scale networks, which will benefit the application of network control methods in the fields of biology and biomedicine. Furthermore NCUA ignores the weight and direction information of network edges, causing a potential bottleneck in characterizing the edge control rather than only node control. Therefore, another important research direction will be the extension of our NCUA method to edge dynamics[31] Taken together, our analysis results support the practical applications of our PNC, which is effective for identifying Personalized driver genes by applying the network control theory to the personalized patient system. Therefore, the PNC reveals new biological insights for clinical applications of personalized therapy and in prognosis analysis.

## Methods

### Paired-SSN

For the Paired-SSN method, the co-expression network of the tumor sample network or normal sample network for each patient is first constructed based on statistical perturbation analysis of one sample against a group of given reference samples (e.g., choosing the expression data of all normal samples of all patients in a given cancer data set as the reference data here) with the SSN method [17]. If the differential PCC (Δ*PCC*) of an edge is statistically significantly large based on the evaluation of the SSN method, this edge would be kept for the single sample (normal sample or tumor sample) from each patient. The p-value for an edge can be obtained from the statistical Z-value measuring Δ*PCC*. Of note, the Δ*PCC* of an edge between genes *i* and *j* and its significant Z-score are calculated using the following formula:

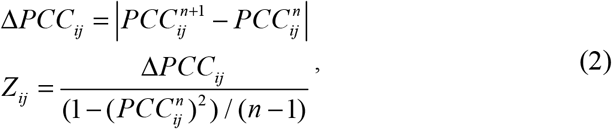

where 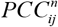 is the PCC of an edge between genes *i* and *j* in the reference network with *n* samples; and 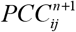is the *PCC* of the edge between genes *i* and *j* in the perturbed network with one additional sample, given that this single sample (e.g. normal sample or tumor sample) for each patient is added to the reference sample group. All of the edges with significantly differential correlations (e.g., p-value < 0.05) are used to constitute the SSN for one normal sample or tumor sample.

Then, the personalized differential co-expression network between the normal sample network and tumor sample network can be constructed in which the edge between gene *i* and gene *j* will exist if the *p*-value of the edge is less than (greater than) 0.05 in the tumor network but greater than (less than) 0.05 in the normal network for their corresponding patient. To more precisely demonstrate the regulatory mechanism of personalized state transition networks, we use the gene interaction network to filter the noise of PCC between gene pairs. Finally the personalized state transition networks are obtained in which edges exist in both the gene interaction network and the differential co-expression network for each patient.

### LIONESS

LIONESS reconstructs the individual specific network in a population of tumor samples as the personalized state transition network for each tumor sample[24]. LIONESS constructs the state transition network by calculating the edge statistical significance between all of the tumor samples and the tumor samples without a given single sample. In order to guarantee a fair comparison with SSN and Paired-SSN, we use PCC in LIONESS to model the sample-specific state transition network. The network specific to one sample in terms of the aggregate networks is then:

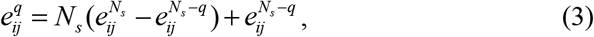

where 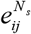 and 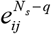 are the PCC values of gene pairs *i* and *j* in sample *n* and all but sample *q*, respectively; *N_s_* is the number of the samples. To construct the sample-specific network, after we obtain the PCC absolute value distribution *S* of all gene pairs we choose a threshold to determine the differential expression edges in the sample-specific network as follows:

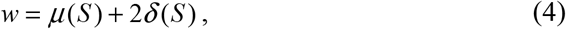

where *μ*(*S*) and *δ*(*S*) are the mean and standard variance of the PCC absolute value distribution *S* of all gene pairs.

### NCUA

In many biological systems, there is adequate knowledge of the underlying wiring diagram, but not of the specific functional forms [32, 33]. Deep analysis of such complicated systems requires structure-based control approaches, which only should know whether there are edges or not. In past decades, the structure based network control studies have shifted from linear dynamics to nonlinear dynamics [34–36]. One of these methods, the FC control method[12, 13] can be reliably applied to large scale complex networks in which the structure is well known and the functional form of the governing equations is not specified. To drive the state of a network to any one of its naturally occurring end states (i.e., dynamical attractors), FC needs to manipulate a set of nodes (i.e., the FVS) that intersects every feedback loop in the network. Under the FC framework, DFVS is proposed to study the structural control problem of directed networks, selecting a minimum set of driver nodes rendering a directed network structural controllable, which has been recently discussed [8]. DFVS assumes that the state of the nodes in a directed network, characterized by source nodes *s_j_*(*t*) and internal nodes *x_i_*(*t*), obeys the following equations:

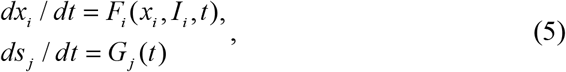

where *I_i_* is a set of predecessor nodes of node *i*, *F_i_* captures the nonlinear response of node *i* to its predecessor nodes *I_i_*, and *G_j_* governs the nonlinear response of source node *s_i_*.

If the undirected networks are considered as bi-directional networks, there are no source nodes in the undirected network nodes. Therefore Eq. (3) of an undirected network *G* (*V, E*), is reduced to,

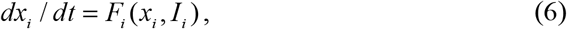

where *x_i_* denotes the state variable of the *i*-th node at time *t* in undirected networks, *I_i_* is a set of neighborhood nodes of node *i*, and *F_i_* captures the nonlinear response of node *i* to its neighborhood nodes in undirected networks.

In fact, the selection of driver nodes may be dependent on the network style (directed or undirected). Motivated by this fact, under the FC framework we formalize the Nonlinear structural Control problem of the Undirected networks (NCU), i.e., how to choose the minimum set of driver nodes for changing the network state from one attractor to the desired attractor in large-scale networks with symmetric edges information. More details can be found in Supplementary Note 1 of Additional File 1. Here we develop a graphic-theoretic algorithm called Nonlinear Control of Undirected network Algorithm (NCUA) under the nonlinear FC framework for determining the minimum driver nodes in undirected networks. Since NCUA and DFVS study the structural network control of undirected and directed biological networks respectively based on the framework of FC, therefore NCUA and DFVS methods completely depict the structure based network control of the large scale system with nonlinear dynamics from the respective of FC.

By assuming that each bi-directional edge is considered as a feedback loop, our NCUA searches the sets of minimum nodes whose removal leaves the graph without edges in undirected networks (Figure 1). The implementation of NCUA consists of two main steps: (i) constructing a bipartite graph from the original undirected network, in which the nodes of the top side are the nodes of the original graph and the nodes of the bottom side are the edges of the original graph, and (ii) determining the MDS of the top side nodes to cover the bottom side nodes in the bipartite graph by using Integer Linear Programming (ILP). The computational details are as follows,

#### Constructing a bipartite graph from the original undirected network

For a given undirected network *G* (*V, E*), we assume that each edge is bi-direct, converting *G* (*V, E*) into a bipartite graph *G* (*V*_T_, *V*_⊥_,*E*_1_), where *V*_T_≡*V* and *V*_⊥_≡*E*. If *v_i_*∈*V*_T_ was one of the nodes for *v_j_*∈*V*_⊥_, we added one edge connecting *v_i_*∈*V*_T_ and *v_j_*∈*V*_⊥_ into the set *E*_1_.

#### Obtaining the cover set with minimum cost by using Integer Linear Programming

After obtaining the bipartite graph *G* (*V*_T_, *V*_⊥_,*E*_1_), we select a minimum dominating set *S* from *V*_T_, which must cover all the nodes in *V*_⊥_. This minimum dominating set is used to determine the driver nodes for controlling the whole network. In NCUA, we select the driver nodes by solving the following ILP:

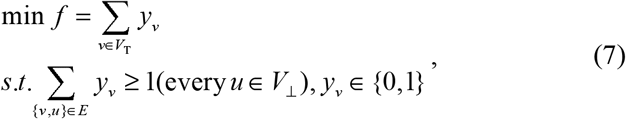

where 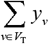 denotes the number of candidate nodes and *y_v_* is an indicative variable; when node *v* in the up side node set in the bipartite network is within the driver nodes, *y_v_*=1, and otherwise, *y_v_*=0; Although it is an NP-hard problem, the optional solution can be obtained for the moderate size graphs with up to a few tens of thousands of nodes by utilizing the LP-based classic branch and bound method [37].

### Hub genes selection method

The Hub genes selection method regards the hub genes in the constructed network as driver genes. After obtaining the degree distribution of all genes *U* in the personalized state transition network, we use a threshold as introduced in following formula to obtain the hub genes,

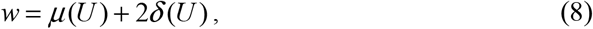

where *μ*(*U*) and *δ*(*U*) are the mean and standard variance of the degree distribution *U* of all gene, respectively.

### F-measure

To verify the effectiveness of the PNC analysis mainly based on structural control methods, the F-measure is considered to assess the enrichment ability of identified personalized driver genes in a given gold standard list (CCG and NCG genes), which considers both the precision and the recall using the formula:

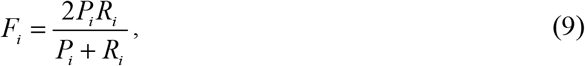

where *P_i_* denotes the fraction of correctly identified genes among all of the identified genes (precision) and *R_i_* denotes the fraction of correctly identified personalized driver genes among the given dataset (recall).

### Enrichment analysis of the personalized driver genes by using hyper geometric test

To estimate the significance of overlap between the predicted personalized driver genes for PNC method and a gold-standard cancer driver gene lists such as Cancer Census Genes and Network of Cancer Genes, we compute the *p*-value by the hyper geometric test [27] as follow,

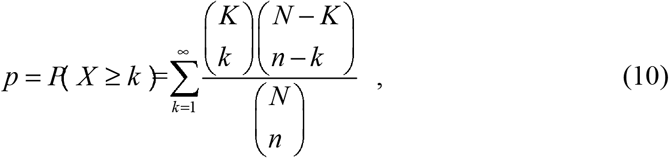

where *N* is the number of genes in background gene interaction network, *K* is the number of gold-standard cancer driver gene lists, *k* is the number of the predicted personalized driver genes overlapping with the genes in the gold-standard cancer driver gene lists, and *n* is number of the predicted personalized driver genes. If the enrichment *p*-value is less than 0.05, then we regard that this predicted personalized driver genes for PNC method is significantly enriched in the gold-standard cancer driver gene lists.

### Availability of code and datasets

The PNC code and the data resources used in this work can be freely downloaded from https://github.com/WilfongGuo/Personalized_Network_Control.

## Appendices

**Additional file 1**: *Additional_file1_Supplemntary_manuscript.docx*. Supplementary manuscript of Structure based control analysis of undirected complex networks with applications to personalized driver genes identification in cancer.

**Additional file 2**: *Additional_file2_CGC_NCG_genes.xlsx*. The list of Cancer Census Genes (CCG) and Network of Cancer Genes (NCG) to assess the F-measure of the identified personalized driver genes of different methods.

**Additional file 3**: *Additional_file3_KS _Personalized_network.xlsx*. The results of personalized network using the Kolmogorov-Smirnov goodness-of-fit test on 13 kinds of cancer datasets from TCGA.

## Authors contributions

WFG and TZ developed the methodology. WFG and YL executed the experiments, WFG wrote this paper. WFG, SWZ, TZ, JG and LC revised the manuscript. SWZ, JG and LC supervised the work, made critical revisions of the paper, and approved the submission of the manuscript. All authors read and approved the final manuscript.

## Acknowledgements

This paper was supported by the National Natural Science Foundation of China (61873202, 61473232, 91430111, 91439103, 91529303, 31771476, 81471047 and 11871456) and National Key R&D Program (2017YFA0505500), Strategic Priority Research Program of the Chinese Academy of Sciences (No. XDB13040700), National Key R&D Program (Special Project on Precision Medicine) (2016YFC0903400), and Natural Science Foundation of Shanghai (17ZR1446100). We thank Professor Tatsuya Akutsu in Kyoto University for giving us valuable comments during the preparation of our manuscript.

