## Additional files for "A novel network control model for identifying personalized driver genes in cancer": Additional_file1_supplementary manuscript1.docx

**Supplementary note 1: The formulation of the of NCU**

Given an undirected network *G* (*V*, *E*), we generally consider the following broader class of model of [1]:

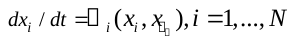
 （S1）

where *x_i_* denotes the state variable of the *i*-th node. The set *I_i_* is a set of neighborhood nodes of node *i*;
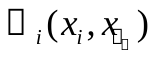
shows the enhancement of activity of node *I_i_*, satisfying that (i) continuous differentiability of
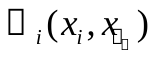
, that is,
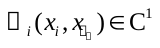
, and (ii) dissipativity, that is, for any initial condition
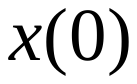
 and for a finite time
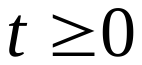
, the dynamical state
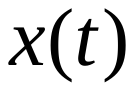
 is bounded by a positive constant *C*:
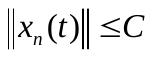
 and iii) decay condition:
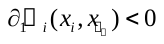
; Similar to the assumption in FVS control in directed networks[2, 3], we formalized the concept of nonlinear structural control of the undirected networks: how we chose the set of input nodes which are injected by input signals
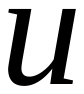
 with the minimum cost to control the system (1) from an initial attractor to a desired attractor. For the system we considered, the following theorem (Theorem 1.3 in [2, 3]) forms the basis of NCU:

*Theorem. Consider a diﬀerential equation system governed by Eq. S1 with dissipative functions*
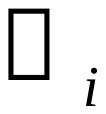
*, and the associated undirected graph G obtained from the I_i_. We also assume*
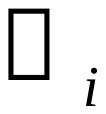
*and its derivatives to be continuous. Moreover, G can contain a self-loop only if Fi does not satisfy the decay condition*
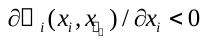
，*Then a nonempty subset J ⊆*
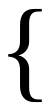
*1,2,...,N*
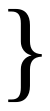
*of vertices of G, and any two solutions X and*
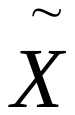
 *of Eq. S1 satisfy*

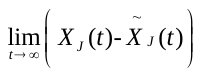
 i*mplies*	(S2)

*for all choices of nonlinearities*
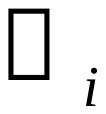
*if and only if J is a feedback vertex set (FVS) of the undirected graph G.*

Basic on the above theorem, we formalize the concept of the Nonlinear Control of the Undirected networks (NCU), which is how we chose proper sets of driver nodes that are injected by input signal with the minimum cost to control the above equation (S1) from an initial attractor to a desired attractor. Note that the minimum FVS in undirected network *G* must exist under our assumption that each edge in undirected network is a feedback loop. Therefore our NCU aims to find minimum FVS in the undirected networks.

**Supplementary note 2:** **supplementary tables**

Table S1: Sample information in TCGA Cancer datasets. Each individual has paired samples (control sample and tumor sample)

| Number | Abbreviation | Description | Number of paired samples |
| --- | --- | --- | --- |
| 1 | BRCA | Breast invasive carcinoma | 112 |
| 2 | COAD | Colon adenocarcinoma | 50 |
| 3 | KICH | Kidney Chromophobe | 23 |
| 4 | KIRC | Kidney renal clear cell carcinoma | 72 |
| 5 | KIRP | Kidney renal papillary cell carcinoma | 31 |
| 6 | LIHC | Liver hepatocellular carcinoma | 50 |
| 7 | LUAD | Lung adenocarcinoma | 57 |
| 8 | LUSC | Lung squamous cell carcinoma | 49 |
| 9 | STAD | Stomach adenocarcinoma | 32 |
| 10 | UCEC | Uterine Corpus Endometrial Carcinoma | 23 |
| 11 | HNSC | Head and Neck Squamous Cell Carcinoma | 43 |
| 12 | PRAD | Prostate Adenocarcinoma | 52 |
| 13 | THCA | Thyroid Papillary Carcinoma | 58 |

Table S2. Concept comparisons between our NCUA and other network control methods.

| Methods | Time Complexity | Network Styles | Dynamics | Targeted state |
| --- | --- | --- | --- | --- |
| MMS | Polynomial time | Directed networks | Local nonlinear | Any |
| MDS | NP-hard | Undirected networks | Nonlinear | Any |
| DFVS | NP-hard | Directed networks | Nonlinear | Attractors |
| NCUA | NP-hard | Undirected networks | Nonlinear | Attractors |

**Supplementary note 3:** **Static model to generate the undirected scale free networks**

We use the following static model [4,5]to generate the undirected network *G* (*V*, *E*) with given exponent
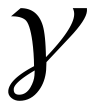
 and the given mean degree
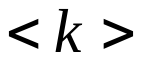
. We start from *N* disconnected nodes indexed by integer number *i* (*i*=1,...*N*). We assign a weight
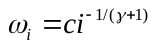
to each node, with a real number in the range [0,1) and *c* is a constant such that

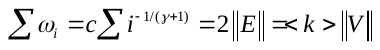
. (S3)

Two nodes *v_i_* and *v_j_* are randomly selected from the set of *N* vertices, with probability proportional to *w_i_* and *w_j_*, respectively. If they have not been connected, then connect them. Otherwise randomly choose another pair. This process is repeated until
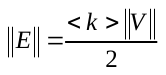
 links are created.

**Supplementary note 4: Fit of power-law distributions on personalized state transition networks by using Kolmogorov-Smirnov goodness-of-fit statistic.**

The personalized state transition networks of different cancer data sets were fitted to power-law distributions to determine the degree exponent γ. First, we computed the connected component (CC) size of each personalized state transition network. The number of nodes in each CC on different cancer data sets is shown in **Additional file 3**. We then used each CC network to fit the power-law distribution. By following [6], let *x* represent a sequence of observations of some variable whose distribution we wish to fit as a power law. There must be some lower bound to the power-law behavior. This value is denoted as *x*_min_. Therefore, given a set containing *n* observations *x_i_* > *x*_min_ and provided that γ>1, it can be shown that the continuous distribution with the corresponding normalizing constant is

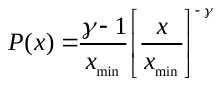
. (S4)

The probability that the data are drawn from a distribution that follows the power law for *x_i_* > *x*_min_ reads as

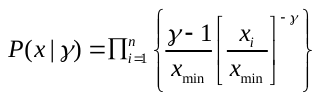
. (S5)

This expression is also called the likelihood of the data given the model. The maximum likelihood estimate (after maximization of the likelihood) for the scaling exponent reads as

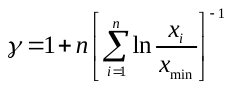
. (S6)

The maximum likelihood, as described above, estimates the scaling parameter γ for each possible value of *x*_min_. The Kolmogorov–Smirnov goodness-of-fit statistic KS is computed. This is done by computing the maximum distance between the cumulative distribution function (CDF) of the data and the fitted model:

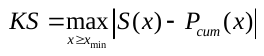
, (S7)

where S(x) is the CDF of the data for the observations with value at least *x*_min_, and *P*_cum_ is the CDF for the power-law distribution that best fits the data. The estimate of *x*_min_ is determined as the value that gives the minimum value KS over all values of *x*_min_. The results for the scaling exponent of personalized state transition networks on different cancer data sets were shown in **Additional file 3**, together with the p-value. The code for calculating the degree exponent γ and the p-value of in Empirical Data are available in http://tuvalu.santafe.edu/~aaronc/powerlaws/.

**Supplementary note 5: Novel controllability findings on synthetic SF network using the NCUA**

In order to evaluate how the network parameters affect control characteristics of undirected networks, we applied our NCUA to the synthetic Scale Free (SF) networks generated by the static model [4,5]. We assumed the degree distribution of the undirected network *G* (*V*, *E*) follows
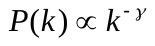
. We first defined the fraction of the driver nodes as the controllability
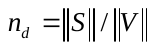
, where
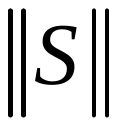
denotes the set of driver nodes to control the whole network and
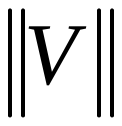
denotes the number of connected nodes in the network. Then, we applied our NCUA to estimate the control characteristics on the synthetic networks. For a given γ and average degree <*k*>, 100 networks of 10,000 nodes were constructed. The fraction of the driver nodes of the NCUA on the synthetic networks was averaged over all realizations. We listed the numerical results of our NCUA for the synthetic networks in **Figure S1**.

In **Figure S1 (a)**, we showed that SF networks with a large value of γ or large value of <*k*> were hard to control, as shown in **Figure S1 (a)**. These results were complemented by **Figure S1 (b-c)**, where it showed that a small fraction of minimum driver nodes were needed to control the entire network if the power law degree exponent γ and average degree <k> was smaller, whereas it was more difficult to controlled with a larger value of γ and average degree <k>. Furthermore when the degree exponent γ is above 3, the minimum number of driver nodes are larger than that of the Erdös-Rényi random networks. This results gave insight into heterogeneous SF networks will be easier to control with the minimum number of driver nodes.

**Figure S1 Controllability of synthetic scale free undirected networks with 10,000 nodes**. All results are averaged over 100 independent realizations of the networks with 10,000 nodes. (**a**) The fraction of driver nodes for NCU control cost *n_d_* in function of the average degree <k> and the degree exponent γ for SF networks. (**b-c**) The fraction of driver nodes for NCU control cost *n_d_* in function of the average degree <k> compared with the ER networks, for the degree exponent γ larger than 2 and less than 2 respectively. It shows that compared with the ER networks, few nodes are needed to control the entire network if the power law degree exponent γ is smaller, whereas more difficultly it is to be controlled, with larger value of γ. These results demonstrate that more heterogeneous the degree distribution of personalized state transition network is, the easier it is to control the entire system.
